# Comparing proportional and ordinal dominance ranks reveals multiple competitive landscapes in an animal society

**DOI:** 10.1101/2020.04.30.065805

**Authors:** Emily J Levy, Matthew N Zipple, Emily McLean, Fernando A Campos, Mauna Dasari, Arielle S Fogel, Mathias Franz, Laurence R Gesquiere, Jacob B Gordon, Laura Grieneisen, Bobby Habig, David J Jansen, Niki H Learn, Chelsea J Weibel, Jeanne Altmann, Susan C Alberts, Elizabeth A Archie

**Author notes:** First authors contributed equally (Emily J Levy, Matthew N Zipple). Last authors contributed equally (Susan C Alberts, Elizabeth A Archie).

## Abstract

Across group-living animals, linear dominance hierarchies lead to disparities in access to resources, health outcomes, and reproductive performance. Studies of how dominance rank affects these outcomes typically employ one of several dominance rank metrics without examining the assumptions each metric makes about its underlying competitive processes. Here we compare the ability of two dominance rank metrics—ordinal rank and proportional or ‘standardized’ rank—to predict 20 distinct traits in a well-studied wild baboon population in Amboseli, Kenya. We propose that ordinal rank best predicts outcomes when competition is density-dependent, while proportional rank best predicts outcomes when competition is density-independent. We found that for 75% (15/20) of the traits, one of the two rank metrics performed better than the other. Strikingly, all male traits were better predicted by ordinal than by proportional rank, while female traits were evenly split between being better predicted by proportional or ordinal rank. Hence, male and female traits are shaped by different competitive regimes: males’ competitive environments are largely driven by density-dependent resource access (e.g., access to estrus females), while females’ competitive environments are shaped by both density-independent resource access (e.g. distributed food resources) and density-dependent resource access. However, traits related to competition for social and mating partners are an exception to this sex-biased pattern: these traits were better predicted by ordinal rank than by proportional rank for both sexes. We argue that this method of comparing how different rank metrics predict traits of interest can be used as a way to distinguish between different competitive processes operating in animal societies.

## Introduction

In group-living animals, individuals can often be linearly ranked according to their priority of access to resources, or their ability to win conflicts (e.g. insects [1–4]; crustaceans [5–7], fish [8–11], birds [12–17], and mammals [18–22]). The resulting dominance hierarchies are associated with a wide range of traits, including physiology [23–25], immunity and disease risk [26–29], behavior [30–32], reproductive success [30,33–38], longevity [30,39–41], and offspring survival [30,35,42,43]. The causes and consequences of dominance rank are therefore integral to our understanding of the evolution of animal behaviors and life history strategies.

When studying these causes and consequences, researchers can choose between several ways of measuring rank (e.g., ordinal rank, proportional rank, Elo score, David’s score [44,45]). Researchers commonly assign each individual’s rank as its order in the dominance hierarchy (i.e., 1, 2, 3, etc.); we refer to this measure as *ordinal rank*. Researchers may also normalize those ordinal ranks to the number of individuals in the hierarchy, producing ranks that represent the proportion of the individuals that each individual dominates (usually referred to as “relative” or “standardized” rank, e.g. [46–54]); we refer to this measure as *proportional rank* because this term describes more precisely the nature of the metric.

Often, researchers choose one of these dominance rank metrics without stating the assumptions that the metric makes about the nature of rank-based competition [46–49,51] (but see [52,55]). The choice of a given rank metric is important because studies sometimes find differences in the ability of different rank metrics to predict rank-related traits, even in the same population. For example, Archie et al. (2014) demonstrated that proportional rank, but not ordinal rank, predicted risk of injury in female baboons in the Amboseli ecosystem in Kenya [56]. In the same population, proportional rank was also a better predictor of females’ fecal glucocorticoid concentrations than ordinal rank (Levy et al., in revision). These studies highlight the need to understand the contexts in which one rank metric predicts a trait better than another.

Here, we examine the ability of ordinal and proportional rank metrics to predict 20 sex- and age-class-specific traits in the Amboseli baboon population (Table 1). We had two goals. First, we explicitly identify the assumptions each metric makes about the underlying competitive landscapes that shape rank-related traits. We identify theoretical scenarios in which we expect either ordinal or proportional rank to be a better measure of competitive interactions and, therefore, a better predictor of rank-related traits. Second, we identify which rank metric (ordinal or proportional) best predicts a wide range of rank-related traits in wild baboons, with the aim of identifying broad patterns of the role of competition in this population.

**Table 1.** Trait descriptions, ΔAICs, and study information for the 20 traits re-analyzed in the present study.

| Trait<sup>a</sup> | Originally Identified Rank Effect<sup>b</sup> | Study Duration (years) | # of Social Groups | Preferred Model<sup>c</sup> | \|ΔAIC\| (Ordinal vs. Proportional) | \|ΔAIC\| (Preferred vs. Null) | Ref<sup>^</sup> |
| --- | --- | --- | --- | --- | --- | --- | --- |
| Percent of consortships obtained (M) | Higher-ranking males obtain more consortships | 12 | 7 | Ordinal | 27 | 158 | [60] |
| Fecal testosterone (M) | Higher-ranking males have higher testosterone levels | 9 | 5 | Ordinal | 24.9 | 27.1 | [61]^ |
| Post-partum amenorrhea duration (F) | Higher-ranking females have shorter periods of post-partum amenorrhea | 36 | 13 | Proportional | 8.5 | 25.9 | [62]^ |
| Inter-birth interval duration (F) | Higher-ranking females have shorter inter-birth intervals | 36 | 13 | Neither | 1.3 | 14.8 | [62]^ |
| Body size (IM) | Juvenile males with higher-ranking mothers have larger body size for their age | 1 | 2 | Ordinal | 7.2 | 15.9 | [63] |
| Wound healing (M) | Higher-ranking males have faster rates of wound healing | 28 | 11 | Ordinal | 8 | 16 | [64] |
| Monthly injury risk (F) | Higher-ranking (proportional) females have a lower risk of injury | 29 | 12 | Proportional | 7.1 | 10.3 | [56]^ |
| Monthly injury risk (M) | Injury incidence is related to a quadratic rank term, with males ranked 3-6 having the highest rates of injury | 28 | 11 | Ordinal | 5.5 | 6.8 | [64] |
| Prenatal fecal estrogen levels (F) | Higher-ranking females have higher prenatal estrogen levels | 1.4 | 5 | Ordinal | 4.2 | 5.8 | [65] |
| Fecal glucocorticoid levels (F)<sup>d</sup> | Higher-ranking females (proportional) have lower glucocorticoid levels | 17 | 12 | Proportional | 3.0 | 1.7 | Levy et al. (in rev)^ |
| Fecal glucocorticoid levels (IM) | Subadult sons of higher-ranking mothers have lower glucocorticoid levels | 4 | 5 | Neither | 0.2 | 4.4 | [66] |
| Fecal glucocorticoid levels (M) | Higher-ranking males have lower glucocorticoid levels (except for the alpha male) | 9 | 5 | Ordinal | 3.8 | 3.6 | [61]^ |
| Time off nipple (IB) | Infants of higher-ranking females tended to spend more time on the nipple | 1.4 | 5 | Ordinal | 3.2 | 6.8 | [65] |
| Initiation rate by infants (IF) | No statistically significant effect of maternal rank on infant initiation rate | 1.4 | 5 | Proportional | 3.4 | 2.5 | [65] |
| Age at menarche (F) | Females with higher-ranking mothers achieve menarche at younger ages | 26 | 9 | Neither | 0.7 | 2.6 | [67] |
| Age at testicular enlargement (M) | Males with higher-ranking mothers achieve testicular enlargement at younger ages | 22 | 9 | Ordinal | 8.5 | 2.3 | [67] |
| Relative infant survival (F) | Higher-ranking females have higher rates of infant survival | 16 | 6 | Neither | 1.2 | 4.0 | [68] |
| Sexual swelling length (F) | Higher-ranking females have longer sexual swellings | 1.5 | 5 | Neither | 0.1 | 2.4 | [69]^ |
| Social connectedness to males (F) | Higher-ranking females are more socially connected to males | 16 | 8 | Ordinal | 3.3 | 33.1 | [70]^ |
| Frequency of received grooming (F) | Higher-ranking females receive more grooming | 2 | 5 | Ordinal | 2.7 | 28.9 | [71]^ |

<sup>a</sup> M = trait measured in adult males; F = trait measured in adult females; B = trait measured in both males and females, no differentiation by sex; I = trait measured in immature individuals, rank measured as maternal rank
<sup>b</sup> Ordinal rank unless otherwise noted
<sup>c</sup> "Neither" if $|\Delta_{\text{ord-prop}}\text{AIC}| < 2$
<sup>e</sup> Indicates that dataset is publicly available on Dryad
<sup>d</sup> $\Delta\text{AIC}$ between Proportional and Null models = 1.7 (close to the 2-unit threshold) and 1% of the 95% CI of proportional rank overlapped with zero, so we included this trait in analysis

### Assumptions of ordinal and proportional rank metrics

As described above, an individual’s ordinal rank reflects the *order* in which an individual appears in a linear dominance hierarchy (i.e., ranks 1, 2, 3…n; Figure 1) [17,57,58]. In contrast, proportional rank accounts for the number of individuals being ranked (i.e., it accounts for hierarchy size) by measuring the *proportion* of other individuals in a hierarchy that an individual outranks (Figure 1) [46–54]. For example, an individual with proportional rank 0.75 outranks 75% of other individuals in its hierarchy. When the number of individuals in the hierarchy does not vary in a given dataset, ordinal and proportional rank are perfectly correlated. However, if the study contains multiple social groups with different hierarchy sizes, or if hierarchy size varies over time, then ordinal and proportional ranks are no longer interchangeable (see Supplementary Materials, Table S1, Figure S2, and Figure S3).

**Figure 1.**
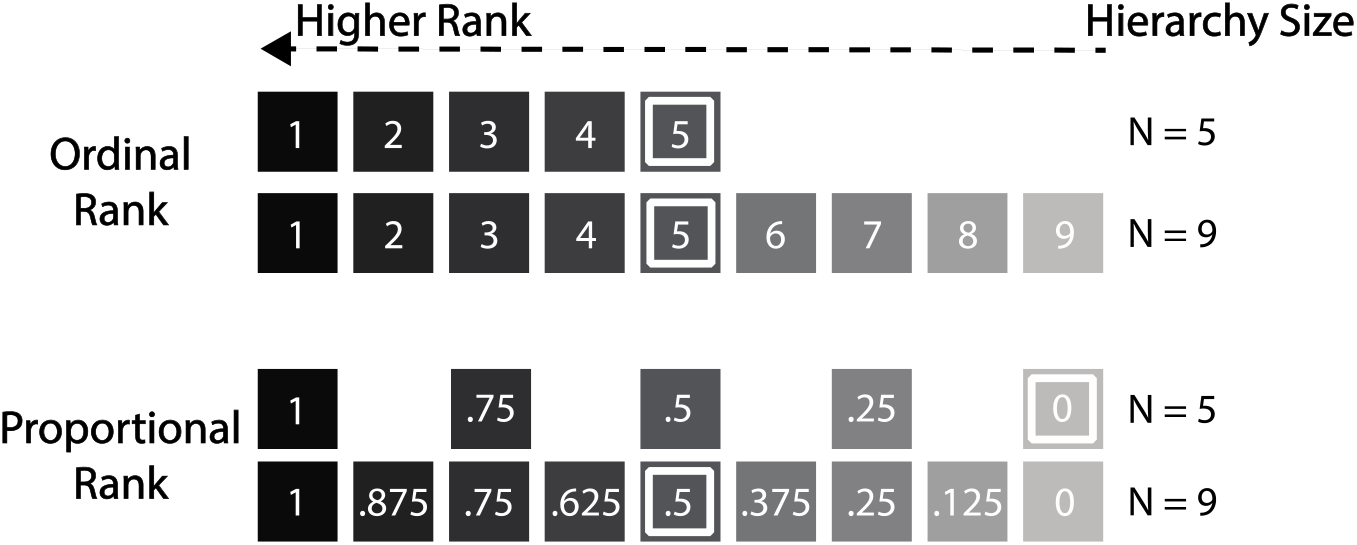
Differences between proportional and ordinal rank in two differently-sized hierarchies. Ranks with darker shading have a competitive advantage over those with lighter shading. The fifth-ranking individual in each hierarchy is demarcated with a white border. Under an ordinal rank framework, being ranked 5^th^ confers the same competitive advantages independent of hierarchy size. Under a proportional rank framework, being ranked 5^th^ is more advantageous in a hierarchy of 9 (proportional rank = 0.5) than in a hierarchy of 5 (proportional rank = 0). Adapted from Levy et al. (in revision).

As a theoretical example of a situation in which ordinal and proportional rank are not interchangeable, consider a hierarchy that contains 5 males. Those males will have ordinal ranks 1-5 and proportional ranks 1, 0.75, 0.5, 0.25, and 0 (Figure 1, N=5). If, over time, four more males join the group and are ranked at the bottom of the hierarchy, the ordinal ranks of the original 5 males will remain the same, but their proportional ranks in the larger hierarchy will be 1, 0.875, 0.75, 0.625, 0.5 (Figure 1, N=9; Figure S3). In this situation, a researcher who uses ordinal rank would conclude that the fifth-ranking male in the hierarchy remained in a constant competitive position throughout the entire study period, whereas a researcher who uses proportional rank would conclude that the fifth-ranking male transitioned from a rank of 0 to 0.5, a major change in dominance rank. Which researcher is correct? The answer depends on the nature of the competitive interactions for which dominance rank serves as a proxy.

The relationship between hierarchy size and resource availability is integral to the assumptions underlying the use of ordinal versus proportional rank metrics (Figure 2; Table S4). Using ordinal rank assumes that the resource base over which individuals compete will not increase as group size increases (Figure 2, orange lines). The result will be more intense competition, on average, in larger groups, and a worse outcome for the lowest-ranking individuals in larger compared to smaller groups. In this scenario, the most salient dominance measure for a focal individual is how many individuals are ranked above that individual. For example, in non-synchronous, non-seasonal breeders such as baboons, only one or two females are likely to be in estrous on any given day, even in large groups – other females may be pregnant, lactating, or in a non-estrous phase of their cycle. This low daily availability of estrous females results in a situation in which the resource over which males compete on a given day (estrous females) increases more slowly than the number of males does, resulting in a decline in average per capita access to estrous females as male hierarchy size increases (Figure 2). If the male dominance hierarchy functions like a queue in which males wait for mating opportunities, a male’s mating opportunities will not depend on the number of other males in his hierarchy per se, but instead upon the number of males that are ranked above him [59] (Table S4, competition for mates). *When average per-capita resource access is density-dependent, we expect ordinal rank to be a better measure of competition and a better predictor of traits determined by that competition compared to proportional rank.*

**Figure 2.**
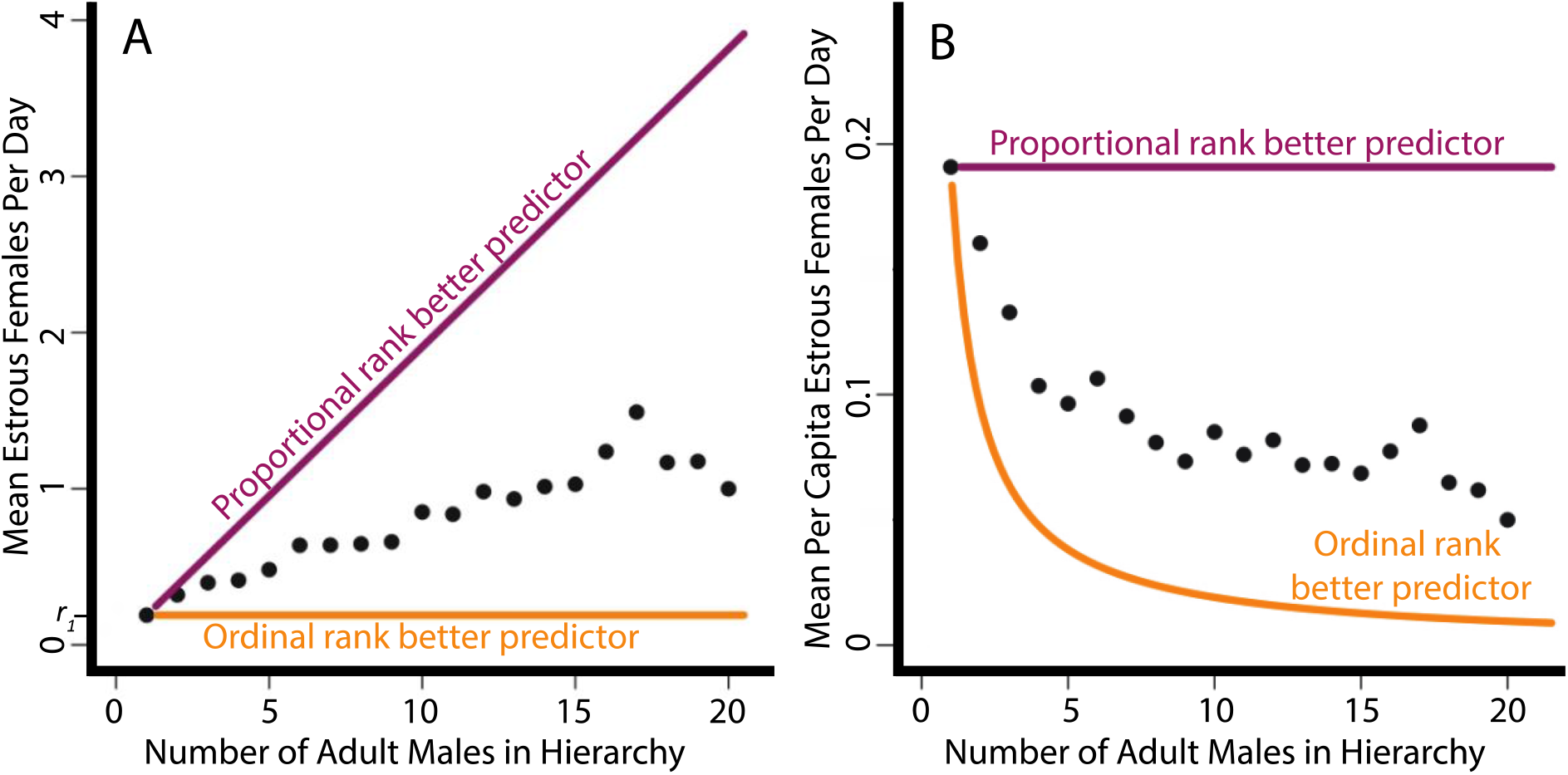
The theoretical and empirical relationships between hierarchy size (x-axis) and resource availability (y-axis), using the example of estrous female baboons, a resource over which male baboons compete for mating success. (A) The orange line shows a theoretical scenario in which the number of estrous females in the group (total resource availability) is constant as the number of males in the hierarchy increases; in this case, male mating success (the resulting measured trait) would be predicted by ordinal rank. The purple line shows a scenario in which the number of estrous females increases in proportion to the number of males in the hierarchy; in this case, male mating success would be predicted by proportional rank. The slope of the orange line is 0 and the intercept is r_1_, which designates the quantity of resources available in a hierarchy size of 1 male (r_1_ = 0.2 estrus females in this figure). This value, r_1_, determines the slope of the purple line; i.e., for proportional rank to perfectly predict mating success, resource availability must increase by r_1_, the quantity available to the first male, as each male is added to the hierarchy. The black points represent empirical data from Amboseli baboons: the empirical relationship between male hierarchy size and the number of estrous females is positive, but the slope is closer to the orange line than the purple line. Thus, we expect ordinal rank to best predict mating success. (B) Similar to (A), but the number of estrous females is plotted per capita (i.e. per adult male in the hierarchy). The orange line illustrates the case in which the resource stays constant across different hierarchy sizes; thus, resources per capita decline as hierarchy size increases. The purple line illustrates the case in which the resource base increases proportionately with hierarchy size; thus, average per capita resource access is fixed. The black points represent the same empirical data as in (A). Note that the framework above assumes that any given individual’s ability to maintain control of a resource is independent of group size.

In contrast, *when average per-capita resource availability is density-independent, such that a larger hierarchy has a proportionately larger resource base, we expect proportional rank to be a better measure of competition and a better predictor of traits determined by that competition compared to ordinal rank* (Figure 2, purple lines). This situation might occur, for instance, in competition for food when a hierarchy of 9 individuals has a home range nearly twice the size (with nearly twice the amount of food) as a hierarchy of 5 individuals. The third-ranking individual in the hierarchy of five has approximately equal access to food as the fifth-ranking individual in the group of nine. In this scenario, the most salient dominance measure for a focal individual is the *proportion* of individuals that it outranks. The individual ranked 5 of 9 is outranked by four individuals, and the individual ranked 3 of 5 is outranked by only two individuals, but both are dominated by 50% of their group mates, and the greater resource base of the larger group means that these two individuals experience approximately the same resource access (Figure 2, purple lines; Table S4, competition for food).

We therefore predict that in any study system in which hierarchy size varies over time or across groups, some rank-related traits will be better predicted by ordinal rank and others will be better predicted by proportional rank. We argue that this difference in predictive power reflects the underlying competitive processes that shape the resulting traits – specifically, the relationship between hierarchy size and resource base. We assess this prediction by examining 20 traits measured as part of a long-term longitudinal study of a wild baboon population, in which both sexes form linear dominance hierarchies. By comparing the differential power of ordinal and proportional rank metrics to predict these 20 traits in this population, we perform the most extensive comparison to date of the ability of different rank metrics to predict traits.

## Results

We identified previous publications from the Amboseli Baboon Research Project that examined relationships between rank and 20 different traits (Table 1). For each trait, we replicated the dataset and statistical model used in the original manuscript, which used either ordinal or proportional rank, by downloading data sets archived in Dryad or by querying the project’s long-term database when archived data were not available. We then built three models for each trait: one using the original rank metric (ordinal or proportional), a second using the alternative rank metric, and a null model that did not include a measure of rank. All models included the same covariates that were used in the original publication’s model (e.g., age, season, reproductive state). We extracted AIC scores for all three models and used an AIC difference of > 2 to indicate that one model was preferred over another, with preference for the model with the smaller AIC value. This 2-unit cutoff is standard practice and approximates a p-value of 0.05 [72,73](see Methods for additional details).

### Rank metrics differ in their ability to predict traits

All 20 traits were better predicted by one or both rank metrics than by the null model (ΔAIC > 2). Furthermore, for 15 of the 20 traits (75%), we found that one of the two rank metrics – ordinal or proportional – performed substantially better than the other in predicting a given trait (ΔAIC > 2; Figure 3, Table 1). In addition, in 7 of these 15 models, only one of the two rank metrics performed better than the null model. This means that for 35% of traits (7 of 20), a relationship between rank and a trait of interest would have been undetected if researchers had chosen the alternative rank metric. For example, male fecal glucocorticoid concentrations were predicted by ordinal rank (ΔAIC to Null = 3.6), but not by proportional rank (ΔAIC to Null = −0.2).

**Figure 3.**
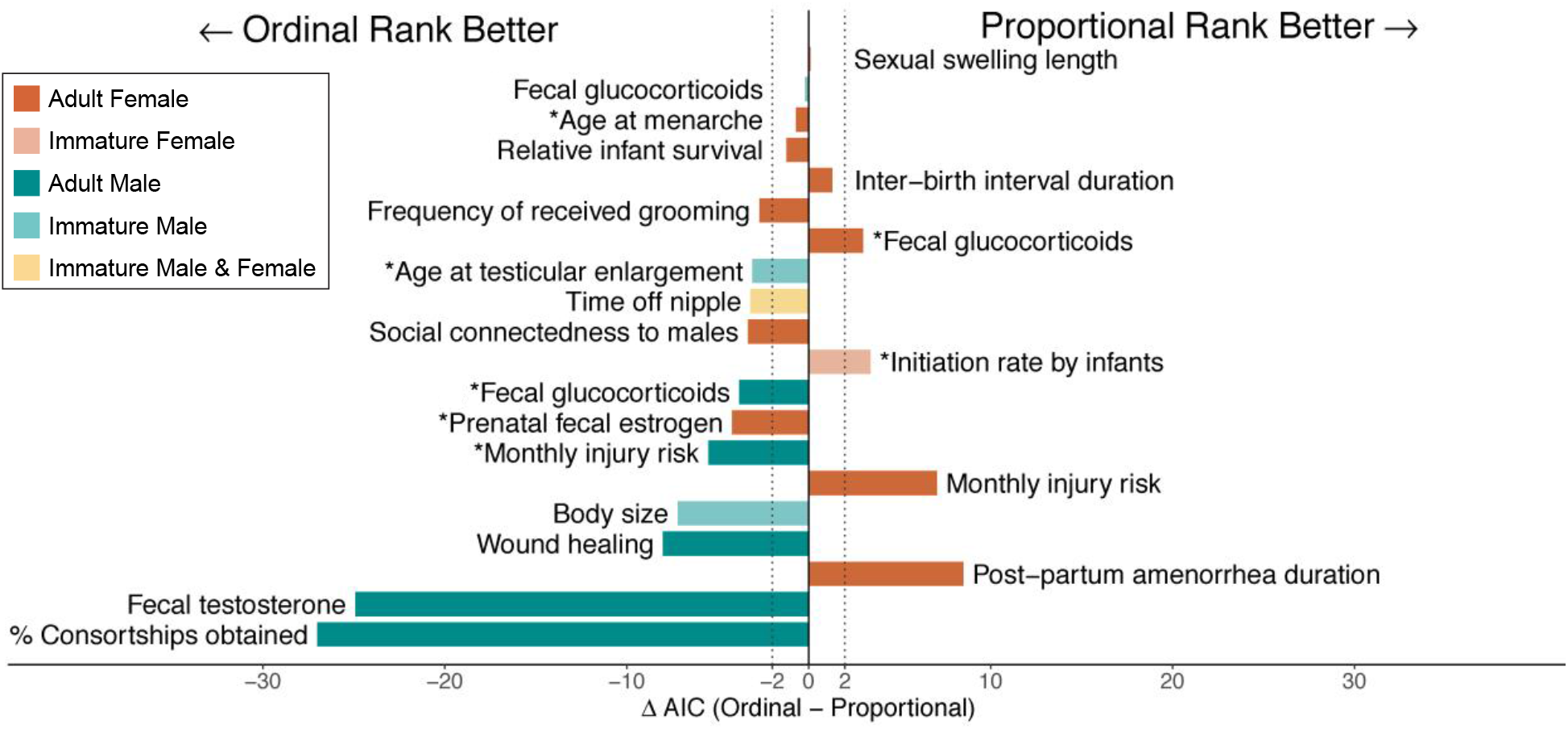
Visualization of model outcomes when predicting the same trait with ordinal versus proportional rank. Each bar corresponds to a trait, and its value corresponds to a difference in AIC scores between models that used ordinal versus proportional rank. Vertical dashed lines represent |ΔAIC| = 2. For traits whose bars are within the dashed lines, neither rank metric performed substantially better than the other (5/20 analyses; we did not find any indication that the ability of the models to differentiate the predictive power of ordinal versus proportional rank depended on the duration of the study; p = 0.9, Pearson’s product moment correlation). For traits whose bars are to the left of the dashed lines, ordinal rank was a better predictor of the trait than proportional rank (11/20), and vice versa for traits whose bars are to the right of the dashed lines (4/20). Colors of bars indicate sex (male, female, both), and shading indicates age class (adult or maternal rank of immatures). Asterisks indicate the seven traits for which only one rank metric predicted the trait better than the null model. The top two bars, sexual swelling length and fecal glucocorticoids, were traits measured in adult females and immature males, respectively.

### All male traits are better predicted by ordinal rank

Whether proportional or ordinal rank was a better predictor of a trait was predicted by the sex of the study individuals, suggesting that male and female baboons experience different competitive regimes. Of the seven male traits that were better predicted by one rank metric than the other, all 7 (100%) were best predicted by ordinal rank (male vs. chance p = 0.02, two-tailed binomial test). In contrast, of the seven female traits that were better predicted by one rank metric than the other, 4 (57%) were best predicted by proportional rank, and 3 (43%) were best predicted by ordinal rank (female vs. chance p = 1.00, two-tailed binomial test; male vs. female p = 0.07, Fisher’s exact test). In two of the three cases where traits could be directly compared between adult males and females (fecal glucocorticoid concentrations and monthly injury risk), male traits were better predicted by ordinal rank whereas female traits were better predicted by proportional rank. Additionally, the trait with the largest AIC difference between rank metrics was percent of consortships obtained by males, which was best predicted by ordinal rank. The ability of male baboons to obtain consortships with females approximates a queuing system [74], such that the most salient feature to a focal male is the number of males ranking higher than him. This pattern is consistent with our understanding of the contexts in which ordinal rank will be a better predictor of resource availability than proportional rank (see *Assumptions of ordinal and proportional rank metrics*, and Figure 2, above).

### All traits related to social and/or mating partners are best predicted by ordinal rank

A second pattern that emerged from these results is that competition for social and mating partners in both sexes was better predicted by ordinal rank than by proportional rank. In all three cases where the trait could be interpreted in terms of access to social partners (male percent consortships obtained, female social connectedness to males, female frequency of received grooming from males or females), the trait was best predicted by ordinal rank. Furthermore, fecal testosterone concentrations in males, which are related to competition for female mating partners [75], was also much better predicted by ordinal rank than by proportional rank.

## Discussion

Ordinal and proportional rank metrics make different assumptions about competitive regimes in animal societies. When average per capita resource access is density-dependent, ordinal rank should predict competition-related traits. In contrast, when average per capita resource access is density-independent, proportional rank should predict competition-related traits. In reality, competition within animal social groups, which experience dynamic, ongoing changes in group size and resources, will rarely be purely density-dependent or density-independent. Instead, most competition will reflect a mix of these two regimes. This point is illustrated in Figure 2 for one resource important to males (number of estrous females); neither density-dependence nor density-independence perfectly describes the relationship between group size and resource availability. Nonetheless, in many contexts, one or the other competitive regime will predominate. In support, we have shown that proportional and ordinal rank metrics differ in how well they predict 75% (15/20) of rank-related traits examined in the Amboseli baboon population. Further, our data indicate that male and female traits tend to be shaped by different competitive regimes: males’ competitive environments appear to be shaped mostly by density-dependent resource access, as evidenced by the strong and consistent performance of the ordinal rank metric in predicting many male phenotypes. In contrast, density-independent competition seems to better describe the competitive regimes that shape many (but not all) female traits. Our results also suggest that, for both sexes, average per-capita access to social and mating partners decreases as hierarchy size increases. Therefore, competition for social and mating partners may be better understood as a density-dependent process.

Because proportional and ordinal ranks reflect different assumptions about the competitive processes influencing social animals, the methods we use here can be applied in other social systems to inform researchers’ understanding of the competitive processes operating in their study species. A researcher who tests proportional and ordinal rank models and finds that ordinal rank is a much stronger predictor of a trait (e.g., male access to females, Figures 2 & 3, Table S4) can conclude that average per-capita access to the resource declines as hierarchy size increases, and that competition for that resource is primarily a density-dependent process. In contrast, a finding that proportional rank better explains a trait (for example, post-partum amenorrhea duration in females, Figure 3), allows a researcher to conclude that the trait is shaped primarily by density-independent competitive processes, such that per-capita access to resources are relatively constant across hierarchy sizes. These methods and logic can also be applied to other rank metrics, such as Elo rating and coding individuals as alpha or non-alpha. Each metric assumes a different underlying competitive process – for example, coding individuals as alpha (highest-ranking) or non-alpha assumes that the alpha individual experiences a different level of resource competition than all others in the hierarchy, who in turn experience comparable resource competition with each other. Models that use each metric can then be compared via AIC score similarly to the present study (Levy et al, in revision).

Our study is the first systematic comparison of the ability of different dominance rank metrics to predict numerous traits in the same population. Proportional and ordinal ranks have rarely been explicitly compared; to our knowledge, only five studies, all in primate species, have previously compared the predictive ability of these two rank metrics. Two studies found that proportional rank better predicted the phenotypes in question than did ordinal rank (male consortship rates in rhesus macaques [55] and rates of injury among female baboons [52]). A third study, in female baboons, found that proportional rank better predicted fecal glucocorticoid concentrations than did ordinal rank, but whether a female had alpha status or not was an even better predictor than proportional rank (Levy et al., in revision). Similarly, a fourth study reported that a ‘high versus low’ categorical measure of rank better predicted female feeding time in rhesus macaques than did proportional or ordinal rank, with high-ranking females spending more time feeding than low-ranking females [76]. A fifth study found that neither proportional nor ordinal rank was a statistically significant predictor of the probability of conception in female blue monkeys [77]. In addition, several methods-based studies have tested whether rank orders differ depending on which metric is used to calculate dominance rank, but none have used empirical data to compare how rank metrics perform in predicting traits (e.g. [78–81]).

Our results also point to the value of long-term, individual-based research [82,83]. Without many years of data or data from multiple social groups, we would have been unable to detect differences in the explanatory power of proportional versus ordinal rank metrics. Through the comparison of these two metrics, we are able to gain a deeper understanding of the sex-specific competitive environments shaping different traits in our study population. We see the previously unappreciated differences in proportional and ordinal rank metrics not as a weakness of research that has already been performed, but as a new tool that can be employed in the study of diverse systems.

Our findings also have implications for meta-analyses and comparative studies of rank-related effects (e.g. [30,32,84]). It is paramount that, before including studies that employ different measures of rank, a meta-analyst considers whether rank metrics presented across multiple studies are equivalent. For example, studies that report effects of rank for “high” versus “low” ranking individuals create category thresholds based on either proportional or ordinal ranks, depending on whether “high” and “low” refers to social position relative to the whole population (ordinal rank) or to each social group individually (proportional rank). Further, if a study is reporting on only a single social group over a short time period, then hierarchy size is likely to be constant and therefore ordinal and proportional ranks would be equivalent. However, if a study is reporting on multiple study groups or even a single study group over a long time period, then rank metrics may no longer be interchangeable. We therefore recommend that meta-analysts assembling datasets from multiple studies should (1) carefully consider the underlying assumptions that link rank metrics to competitive landscapes in order to determine which rank metric is most appropriate, and (2) include only studies with equivalent rank metrics in a given meta-analysis, converting between rank metrics when possible and necessary. When following these recommendations is impracticable, meta-analysts should acknowledge the limitations of drawing inferences from studies with non-equivalent rank metrics.

We hope that our findings encourage other researchers working on long-term studies to perform similar analyses comparing the predictive power of proportional and ordinal rank metrics. We also encourage researchers to consider and explicitly state the latent assumptions that are made by using any particular rank metric and to consider if their traits of study are more likely to be explained by one rank metric versus another.

## Acknowledgements

This project would not have been possible without long-term collaborators who have helped us maintain this project for nearly 50 years. We gratefully acknowledge the support of the National Science Foundation and the National Institutes of Health for the majority of the data represented here, currently through NSF IOS 1456832, and through NIH R01AG053308, R01AG053330, R01HD088558, and P01AG031719. We also thank Duke University, Princeton University, and the University of Notre Dame for financial and logistical support. We also thank the Kenya Wildlife Service, University of Nairobi, Kenya Institute of Primate Research, National Museums of Kenya, the Kenya National Council for Science, Technology and Innovation, and the Enduimet Wildlife Management Association. We also thank the members of the Amboseli-Longido pastoralist communities, Ker & Downey Safaris, Air Kenya, and Safarilink for their cooperation and assistance in the field. Particular thanks go to the Amboseli Baboon Research Project long-term field team – R.S. Mututua, S.N. Sayialel, J.K. Warutere, and I.L. Siodi - and to T. Wango and V. Oudu for their untiring assistance in Nairobi. The baboon project database, Babase, is designed and programmed by K. Pinc. This research was approved by the IACUC at Duke University, University of Notre Dame, and Princeton University and adhered to all the laws and guidelines of Kenya. For a complete set of acknowledgments of funding sources, logistical assistance, and data collection and management, please visit http://amboselibaboons.nd.edu/acknowledgements/.

## Methods

### Study Population

The Amboseli Baboon Research Project is a long-term study of a natural population of savannah baboons located in Kenya’s Amboseli basin. Data collection began in 1971 and continues today [85]. The population consists primarily of yellow baboons (*Papio cynocephalus*) that experience some naturally-occurring admixture with olive baboons (*P. anubis*) [86–88]. The number of social groups under observation at any given time has ranged from 1 to 6, varying either as a result of logistical considerations, group fissions, or group fusions. All individuals in study groups are visually recognized based on morphological and facial features. Near-daily demographic, environmental, and behavioral data have been collected throughout the study, and paternity data (beginning ca. 1995) and endocrinological data (beginning ca. 2000) have been collected for part of the study.

### Calculation of Dominance Rank

Dominance ranks are determined by assigning wins and losses in dyadic agonistic interactions between same-sex individuals. Data on agonistic interactions are collected ad libitum during daily data collection, typically while the observer is simultaneously carrying out random-order focal animal sampling [89]. This sampling procedure ensures that observers continually move to new locations within the social group and observe focal individuals on a regular rotating basis. An individual is considered to win an agonistic interaction if they displace another individual, or if they give only aggressive or neutral gestures while their opponent gives only submissive gestures. All agonistic outcomes are entered into sex-specific dominance matrices (i.e., males are ranked separately from females). Individuals are placed in order of descending, sex-specific rank so as to minimize the number of entries that fall below the diagonals of the matrices [90,91]. Ranks are assigned for all group members every month. Only adult ranks are considered in this analysis.

Ordinal ranks are produced by numbering individuals according to the order in which they occur on the monthly matrix (1,2,3…n, where n = hierarchy size), with 1 being the highest-ranking male or female in the hierarchy and *n* being the lowest. Proportional ranks are computed as (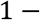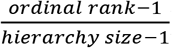) to produce ranks that fall in a range of [0,1] in every hierarchy, with 1 still being the highest-ranking male or female in the hierarchy and 0 being the lowest.

### Re-analysis of Previous Studies

We aimed to test whether 20 different sex- and age-class-specific traits were better predicted by ordinal or proportional rank in the Amboseli baboon population. We first identified previous publications from the Amboseli project that reported statistically significant effects of rank on various traits. For a complete list of re-analyses performed, see Table 1.

Our methods of re-analysis followed three steps:

1. We replicated as closely as possible the dataset used to produce the original analyses. In the case of datasets stored on the Dryad Digital Repository (datadryad.org), these datasets could be matched exactly (see Table 1). If the original dataset was not deposited on Dryad, we re-extracted the dataset as well as we could from the Amboseli Baboon Research Project’s long-term, relational database. However, the datasets we extracted were sometimes slightly different from those originally analyzed, because the database changes slightly over time as corrections are made. In all cases, we produced qualitatively close matches to the originally reported dataset in terms of sample sizes and summary statistics.
2. We replicated as closely as possible the models presented in the original analysis. All re-analyses were carried out in R [92], even though some original analyses were carried out in SPSS, JMP, or SAS. In order to maintain consistency across all analyses reported here, all linear models, general linear models, and mixed effects models were built using the function *glmmTMB* [93]. All survival models were built using the function *coxph* [94]. In some cases, differences between the original study and our replication, either because of software differences or dataset differences, caused our replicated models to be slightly different from the original models. However, our re-analyses were qualitatively consistent with the original analyses.
3. For each of the models described in step 2, we built two additional alternative models: (1) A model that replaced the rank term used in the original model with the alternative rank metric (proportional rank if ordinal rank was originally used and vice versa). (2) A null model that removed the rank term from the model. We then extracted AIC values from all three models to determine which model, if any, best fit the data. We interpreted an AIC difference of ≥ 2 to mean that one model was preferred over another, with preference for the model with a lower AIC score.

## Supplementary Materials

### Identifying changes in the relationship between ordinal and proportional ranks over time

In long-term studies, hierarchy size varies over time and across social groups. This variation should simultaneously weaken the relationship between ordinal and proportional rank and increase our ability to measure different competitive processes in social groups. To test the prediction that the relationship between ordinal and proportional ranks weakens as studies progress, we measured the correlation between monthly ordinal and proportional ranks in the Amboseli Baboon Research Project dataset over increasingly longer periods of time.

Specifically, for each social group we have studied (N = 17 groups), we calculated the R^2^ values from linear models that predicted proportional rank as a function of ordinal rank using increasingly larger datasets. The decision of which metric to use as the predictor variable, in this case ordinal rank, and which as the response variable, in this case proportional rank, was random and had no effect on the results of these analyses. We began by calculating this correlation using only rank data from the first month that a group was under observation (R^2^ necessarily = 1). We then repeated this R^2^ calculation iteratively, each time drawing on ever-larger datasets, by adding data in 12-month increments (i.e. 13 total months, 25 total months, 37 total months, etc.), until we reached the last available dataset of ranks for a group (see Table S1 an example dataset). This method allowed us to track the strength of the predictive relationship between ordinal and proportional ranks as the study progressed.

These analyses included a total of 17 social groups that have been studied over the last 40+ years (thin black lines in Figure S2). We also repeated the same approach, combining data from all social groups into a single analysis (thick grey lines in Figure S2), allowing us to qualitatively compare patterns of change in the relationship between ordinal and proportional ranks both within social groups and across the study population. Note that at the start of the project, only a single social group was followed (Alto’s group). As a result, the grey line starts at an R^2^ value of 1. If multiple study groups with different group sizes had been followed at the beginning of the study, the R^2^ value at the beginning of the project would have been < 1.

**Table S1.** Example dominance ranks from seven individuals across three months and how these data would be used to calculate R^2^ values via a linear model for predicting proportional rank from ordinal rank. To calculate the relationship between ordinal and proportional ranks across the 3 months in the table (January, February, and March 2016), every row in this dataset would be used in a linear model in which proportional rank is the response variable and ordinal rank is the predictor variable. Individual identity did not factor into the model or calculation of R^2^, so the switch in rank order between individuals C, D, and E from February to March 2016 is irrelevant. What does, however, reduce R^2^ is the addition of individual G to the group in February 2016, and the loss of individual G from the group in March 2016.

| Individual Identity | Year-Month | Ordinal Rank | Proportional Rank |
| --- | --- | --- | --- |
| A | Jan-2016 | 1 | 1 |
| B | Jan-2016 | 2 | 0.8 |
| C | Jan-2016 | 3 | 0.6 |
| D | Jan-2016 | 4 | 0.4 |
| E | Jan-2016 | 5 | 0.2 |
| F | Jan-2016 | 6 | 0 |
| A | Feb-2016 | 1 | 1 |
| B | Feb-2016 | 2 | 0.8333 |
| C | Feb-2016 | 3 | 0.6667 |
| D | Feb-2016 | 4 | 0.5 |
| E | Feb-2016 | 5 | 0.3333 |
| F | Feb-2016 | 6 | 0.1667 |
| G | Feb-2016 | 7 | 0 |
| A | Mar-2016 | 1 | 1 |
| B | Mar-2016 | 2 | 0.8 |
| E | Mar-2016 | 5 | 0.2 |
| C | Mar-2016 | 4 | 0.4 |
| D | Mar-2016 | 3 | 0.6 |
| F | Mar-2016 | 6 | 0 |

As predicted, the correlation between ordinal and proportional ranks both within and across study groups decreased as the length of study increased (Figure S2). This is because the size of adult female and male hierarchies changed over time. Variation in hierarchy size, in turn, decouples our density-independent rank metric (proportional) from our density-dependent rank metric (ordinal). This decline in R^2^ as the length of study increased was seen in each group individually and across all study groups when all data were combined (i.e., across the study population).

The decline in R^2^ over time was apparent in both male and female ranks, although the decline occurred more quickly and less linearly in the male rank data than in the female rank data. This sex difference, which we did not predict, prompted us to form two post-hoc predictions to explain it. (1) Baboon groups contained fewer adult males than females; hence hierarchy sizes are smaller for males than for females. The addition of one individual to a small hierarchy changes all members’ proportional ranks more than the addition of one individual to a large hierarchy (Figure S3). Thus, if we assume that different-sized groups have comparable rates of maturation, death, and dispersal, the relationship between ordinal and proportional ranks would be weaker in smaller hierarchies than larger hierarchies. (2) Changes in male hierarchy size were more common than changes in female hierarchy size due to frequent male dispersal, and all changes in hierarchy size reduce the relationship between ordinal and proportional ranks. Together, we would expect these two sex differences – in average hierarchy size and in the frequency of changes in hierarchy size – to lead to differences in the relationship between ordinal and proportional rank between males and females (Figure S3).

We performed post-hoc analyses and confirmed both of our predictions. Of the 1,637 group-months for which we had rank data for both males and females, adult males outnumbered adult females in only 14 months (<1% of group-months; mean # of females – mean # of males ± SD =7.4 ± 0.1, p<0.0001 in one-sample, two-tailed t-test). Additionally, on average, the number of adult males in a social group changed more from one month to the next as compared to the number of adult females (mean absolute change in # adult males per month ± SD = 0.59 ± 0.02, mean in adult females ± SD = 0.25 ± 0.01, p<0.0001 in two-sample, two-tailed t-test).

**Figure S2.**
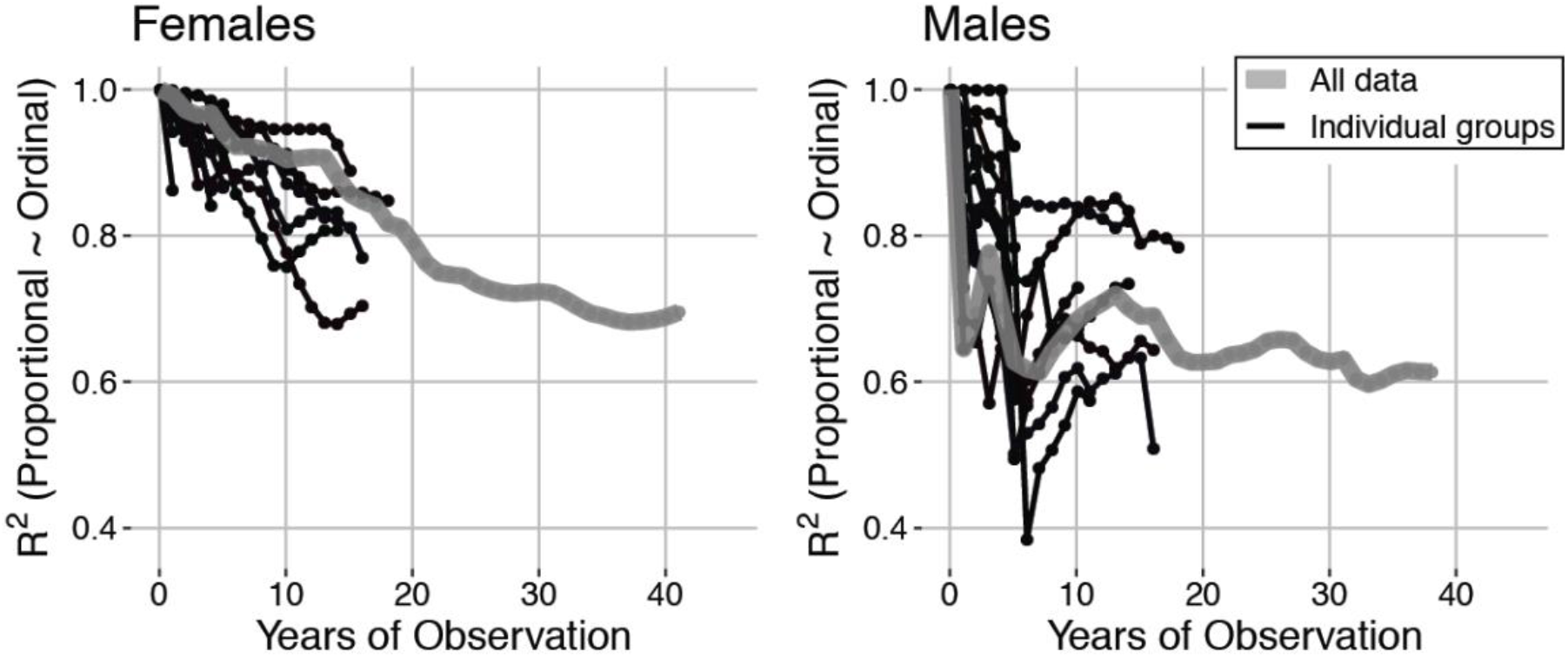
The relationship between ordinal and proportional ranks weakens as the period of observation increases in both females and males. Black points and lines indicate changes in R^2^ for every social group over time, and the grey points and lines indicate changes across all social groups (i.e. across the study population) using pooled data from all individuals in all social groups. At each point, R^2^ was calculated from the cumulative dataset (e.g. the grey point at 13 months includes data from all 13 months across all individuals in all social groups). The grey line extends farther than any black line because the black lines represent individual social groups, which are not permanent due to fissions and fusions, whereas the grey line represents the full dataset.

## Supplementary Figures and Tables

**Figure S3.**
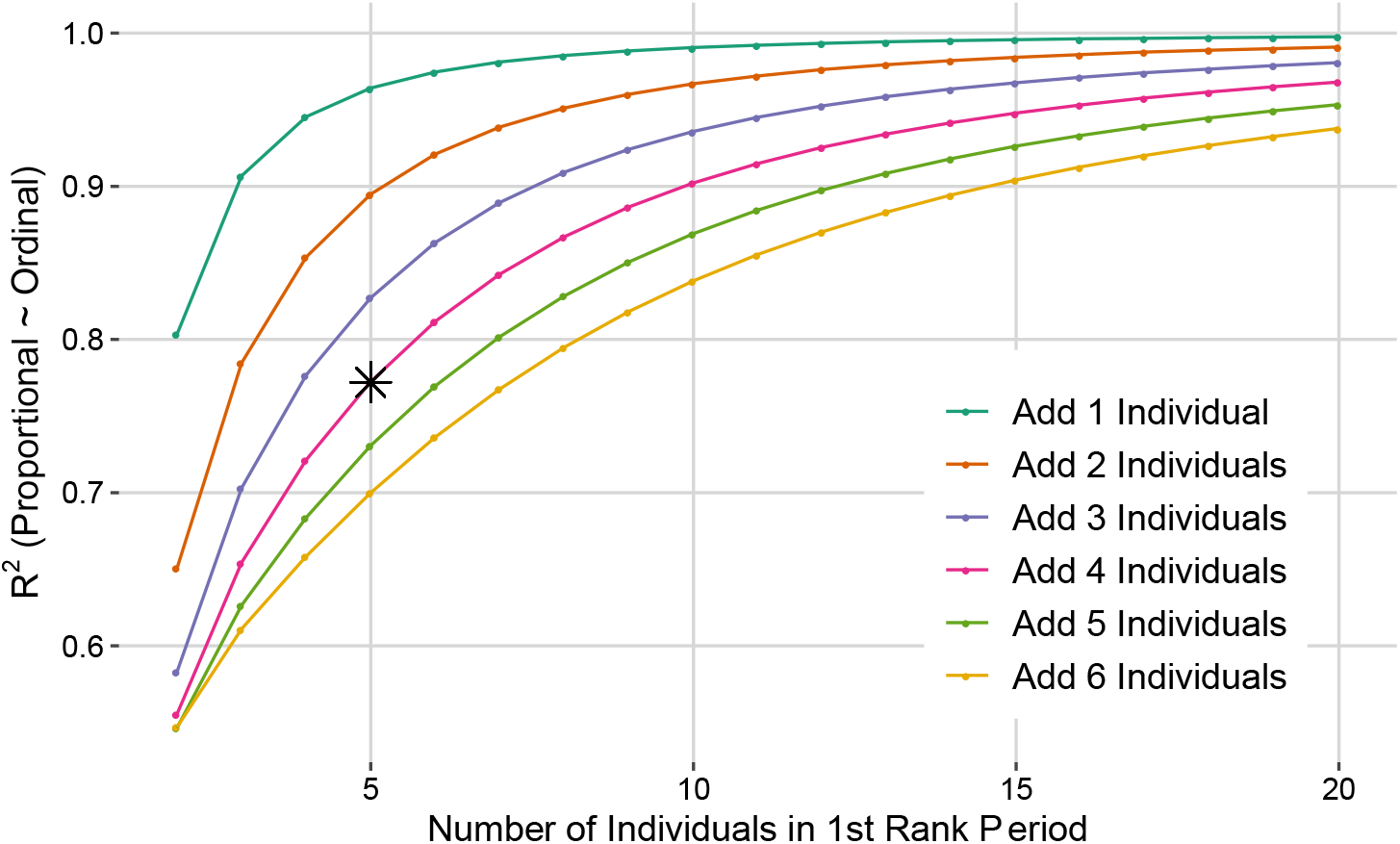
The relationship between ordinal and proportional ranks is weakened when hierarchy size changes. We simulated groups that included a varying numbers of adults (range 2-20) and assigned ordinal and proportional ranks to each individual for one rank period (one month). We then added a varying number of individuals to the group, and again assigned ordinal and proportional ranks to each individual for a second ranking period. We then calculated the R^2^ value from a model that predicted proportional rank as a function of ordinal rank, including all ranks from both ranking periods. The relationship between ordinal and proportional ranks is less robust to greater changes in hierarchy size and less robust to changes in smaller starting hierarchies. The situation described in the introduction, in which four males join an existing group of five males, is marked with an asterisk.

**Table S4.**
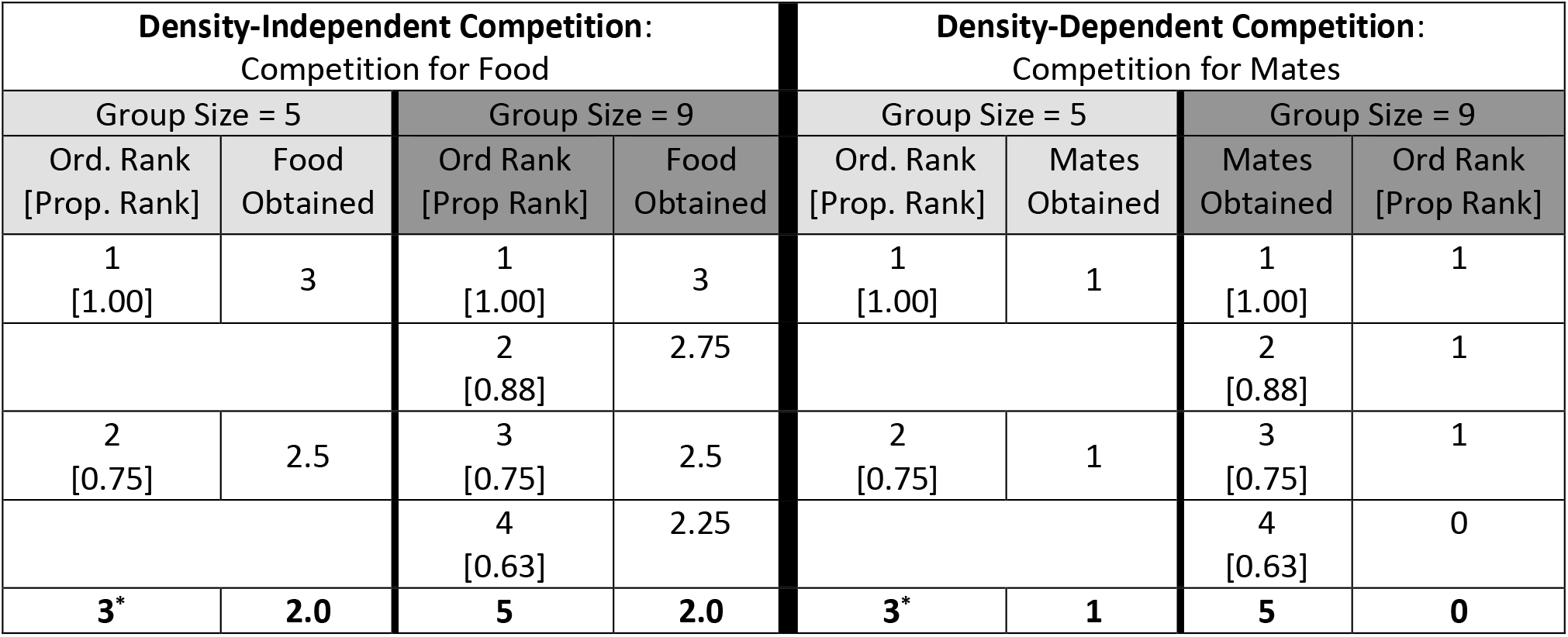
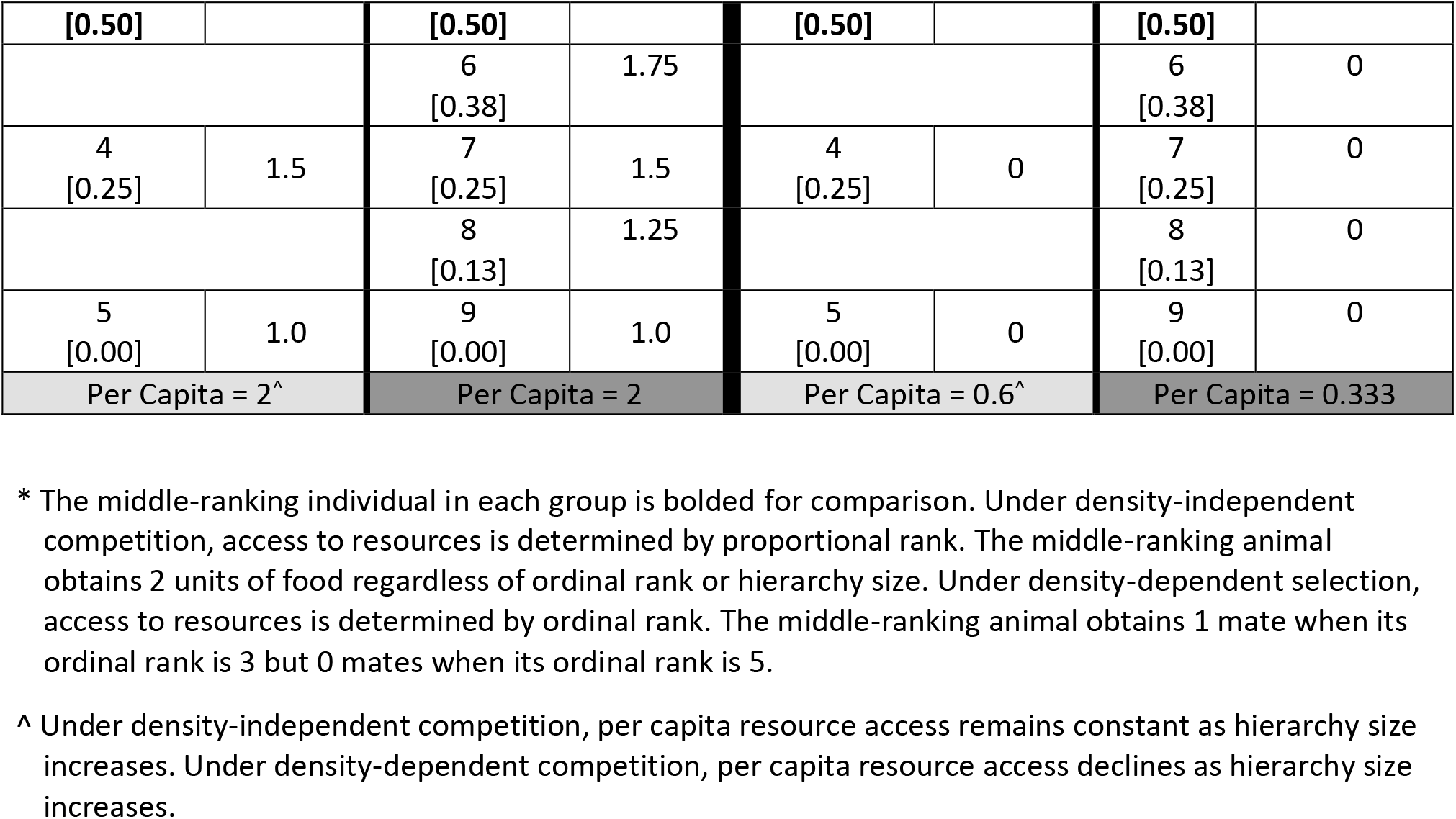
Theoretical distribution of resources under density-independent and density-dependent competition for two group sizes to demonstrate theoretical differences between ordinal and proportional ranks.

| Density-Independent Competition:<br>Competition for Food |  |  |  | Density-Dependent Competition:<br>Competition for Mates |  |  |  |
| --- | --- | --- | --- | --- | --- | --- | --- |
| Group Size = 5 |  | Group Size = 9 |  | Group Size = 5 |  | Group Size = 9 |  |
| Ord. Rank<br>[Prop. Rank] | Food<br>Obtained | Ord Rank<br>[Prop Rank] | Food<br>Obtained | Ord. Rank<br>[Prop. Rank] | Mates<br>Obtained | Mates<br>Obtained | Ord Rank<br>[Prop Rank] |
| 1<br>[1.00] | 3 | 1<br>[1.00] | 3 | 1<br>[1.00] | 1 | 1<br>[1.00] | 1 |
|  |  | 2<br>[0.88] | 2.75 |  |  | 2<br>[0.88] | 1 |
| 2<br>[0.75] | 2.5 | 3<br>[0.75] | 2.5 | 2<br>[0.75] | 1 | 3<br>[0.75] | 1 |
|  |  | 4<br>[0.63] | 2.25 |  |  | 4<br>[0.63] | 0 |
| **3\*** | **2.0** | **5** | **2.0** | **3\*** | **1** | **5** | **0** |
| **[0.50]** |  | **[0.50]** |  | **[0.50]** |  | **[0.50]** |  |
|  |  | 6<br>[0.38] | 1.75 |  |  | 6<br>[0.38] | 0 |
| 4<br>[0.25] | 1.5 | 7<br>[0.25] | 1.5 | 4<br>[0.25] | 0 | 7<br>[0.25] | 0 |
|  |  | 8<br>[0.13] | 1.25 |  |  | 8<br>[0.13] | 0 |
| 5<br>[0.00] | 1.0 | 9<br>[0.00] | 1.0 | 5<br>[0.00] | 0 | 9<br>[0.00] | 0 |
| Per Capita = 2<sup>^</sup> |  | Per Capita = 2 |  | Per Capita = 0.6<sup>^</sup> |  | Per Capita = 0.333 |  |

\* The middle-ranking individual in each group is bolded for comparison. Under density-independent competition, access to resources is determined by proportional rank. The middle-ranking animal obtains 2 units of food regardless of ordinal rank or hierarchy size. Under density-dependent selection, access to resources is determined by ordinal rank. The middle-ranking animal obtains 1 mate when its ordinal rank is 3 but 0 mates when its ordinal rank is 5.
<sup>^</sup> Under density-independent competition, per capita resource access remains constant as hierarchy size increases. Under density-dependent competition, per capita resource access declines as hierarchy size increases.

